# Single cell response landscape of graded Nodal signaling in zebrafish explants

**DOI:** 10.1101/2021.04.25.441305

**Authors:** Tao Cheng, Yan-Yi Xing, Yun-Fei Li, Cong Liu, Ying Huang, Ying-Jie Zhang, Sean G. Megason, Peng-Fei Xu

## Abstract

Nodal, as a morphogen, plays important roles in cell fate decision, pattern formation and organizer function. But because of the complex context *in vivo* and technology limitations, systematic studying of genes, cell types and patterns induced by Nodal alone is still missing. Here, by using a relatively simplified model, the zebrafish blastula animal pole explant avoiding additional instructive signals and prepatterns, we constructed a single cell response landscape of graded Nodal signaling, identified 105 Nodal immediate targets and depicted their expression patterns. Our results show that Nodal signaling is sufficient to induce anterior-posterior patterned axial mesoderm and head structure. Surprisingly, the endoderm induced by Nodal alone is mainly the anterior endoderm which gives rise to the pharyngeal pouch only, but not internal organs. Among the 105 Nodal targets, we identified 14 genes carrying varying levels of axis induction capability. Overall, our work provides new insights for understanding of the Nodal function and a valuable resource for future studies of patterning and morphogenesis induced by it.

## Introduction

During embryonic development, morphogen gradients play important roles in positional patterning and fate decisions of responding cells (Gurdon & Bourillot, 2001; Rogers & Müller, 2019; Rogers & Schier, 2011; Shilo & Barkai, 2017; Soh *et al*., 2020). Many well-researched processes have been shown to be patterned by morphogens, such as germ layer specification (Feldman *et al*., 1998; Kiecker *et al*., 2016), body axes induction (Schier & Talbot, 2005) and organ formation such as the dorsal-ventral patterning of neural tube (Wilson & Maden, 2005). Our previous work has shown that two opposing gradients of Nodal and BMP can lead to the formation of a complete body axis both *in vivo* and in animal pole explants of zebrafish blastulae (Xu *et al*., 2014). A key question raised by this study is what are the individual contributions of Nodal in this robust induction process?

Nodal, as an important member of TGF-beta family, has been shown to induce mesendoderm formation (Dougan *et al*., 2003; Hagos & Dougan, 2007) and pattern the anterior-posterior axis of the embryo in a concentration-dependent manner (Thisse *et al*., 2000). Previous work has provided insights into Nodal function, as well as the formation and interpretation of its gradient (Chen & Schier, 2001; Dubrulle *et al*., 2015; Müller *et al*., 2012; Wang *et al*., 2016). However, an inevitable problem of *in vivo* studies is that it is difficult to untangle the reciprocal regulations between different morphogens intermingled spatiotemporally during development (Zinski *et al*., 2018). At the same time, for master regulators such as Nodal, loss of function studies help illuminate its initial functions during development but not at later developmental stages.

Zebrafish blastula animal pole explants have emerged as a model system that can response to external signals (Ogata *et al*., 2007; Xu *et al*., 2014). Several recent works have used different portions of the zebrafish blastoderm to study the self-organization of naïve cells and morphogenesis (Fulton *et al*., 2020; Schauer & Pinheiro, 2020; Williams & Solnica-Krezel, 2020). We have previously shown that the animal pole half of zebrafish blastoderm contains limited signals and that upon application of the appropriate signals, it is able to recapitulate normal developmental processes leading to the formation of an embryonic axis (Xu *et al*., 2014). Therefore, it appears to be an ideal *ex vivo* model for the functional dissection of a single morphogen, as well as studying the interaction between different tissues or germ layers, which is very difficult to achieve *in vivo* or in a 2-D cell culture system.

In this study, using the zebrafish animal pole explants, in combination with comprehensive transcriptome analysis, we identified the first wave of Nodal response genes and established a response landscape of graded Nodal signaling at single cell resolution. Our results show that Nodal signaling by itself is sufficient to induce anterior-posterior patterned dorsal axis mesoderm derivatives and part of the endoderm cells. Furthermore, we identify that the organizer function of Nodal is achieved by its multiple downstream targets which carry various axis induction capabilities.

## Results

### Using zebrafish animal pole explants to study germ layer induction and morphogenesis in response to Nodal signaling

Explants, as a reproducible and controlled *ex vivo* system, have long been used to study developmental signaling and morphogenesis (Ogata *et al*., 2007; Sagerström *et al*., 1996). To better understand Nodal induced cell fate specification and morphogenesis, we established an explant system with an artificial Nodal gradient.

We injected Nodal related 2 (Ndr2 also known as Cyclops) mRNA into a single animal pole blastomere of a zebrafish embryo at 64-128 cell stage (2 – 2.25 hpf (hours post fertilization)). Explants of the animal pole were performed at the 512-cell stage and the formation of the Nodal signaling gradient is assessed by immunofluorescence for phosphorylated Smad2/3 (p-Smad2/3), which are the factors responsible for the transduction of Nodal signals into the nucleus (Supplementary Fig. S1D, Fig. 1C and Supplementary Movie 3-5). Taking explants at the 512-cell stage prevents the stimulation of cells of the explant by inductive signals secreted by the embryonic margin (Xu *et al*., 2014).

**Figure 1.**
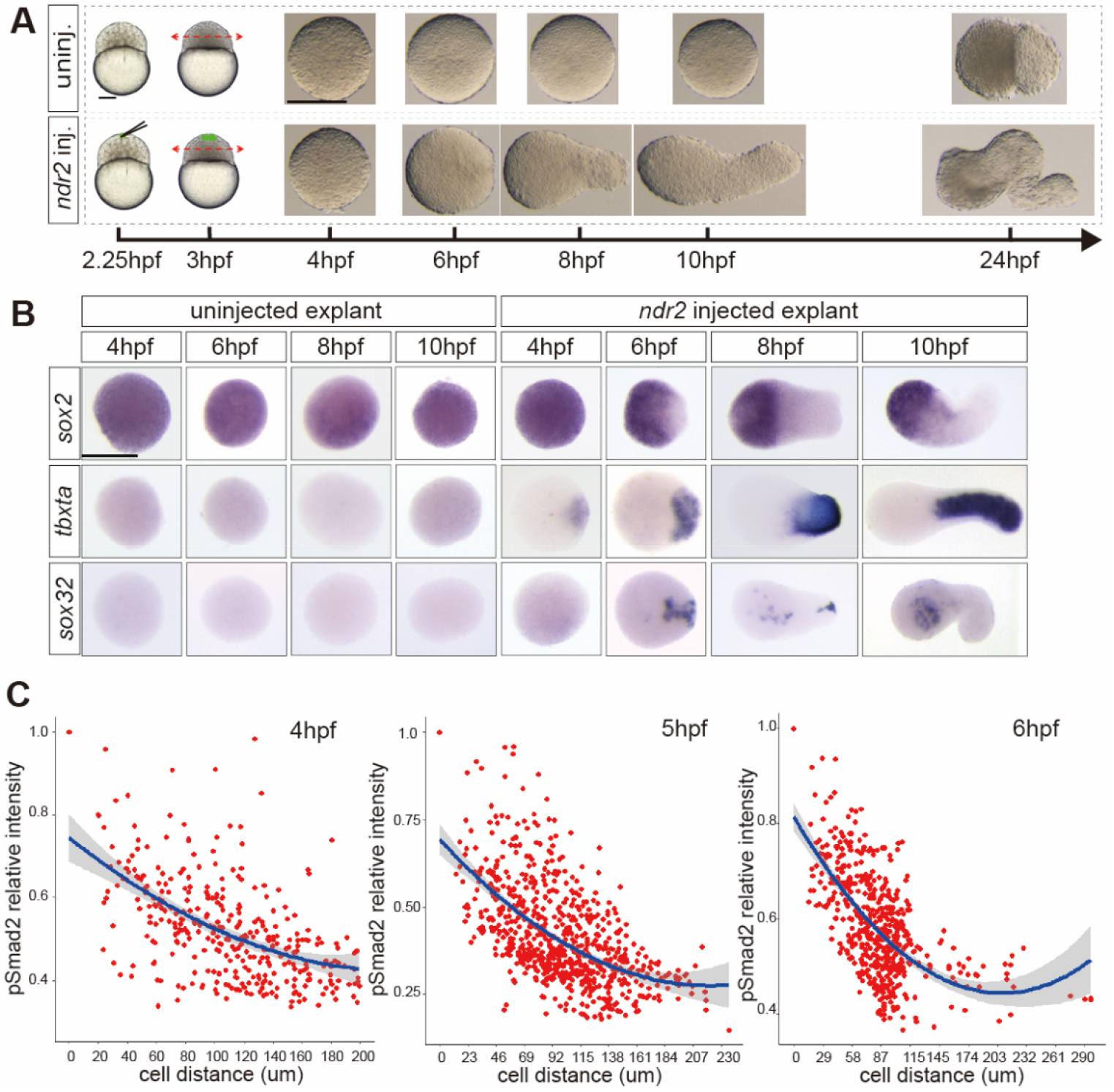
Morphogenesis, germ layers and patterning induced by Nodal gradient in zebrafish explants. **A,** One blastomere at the animal pole was injected with 10 pg *ndr2* mRNA at 64-128 cell stage, uninjected embryos are used as control. The animal half of the injected and uninjected blastoderm were cut off at 3 hpf and cultured in serum replacement medium. Nodal injected explants showed obvious elongation from 6 hpf, while uninjected explants (upper) did not. **B,** Expression patterns of ectoderm (*sox2*), mesoderm (*tbxta*) and endoderm (*sox32*) markers were determined by WISH in both Nodal injected and control explants at indicated stages. **C,** Quantification of P-Smad2/3 gradient in Nodal injected explants at indicated stages. hpf, hours post fertilization. Scale bar: 200 μm (**A**, **B**).

Following explant, both uninjected animal pole explants and Nodal injected explants (referred to as Nodal explants hereafter) initially rounded up. Afterward, the overall morphology of uninjected explants stayed roughly unchanged, while Nodal explants undergo obvious morphogenesis (Fig. 1A and Supplementary Movie 1-2). First, *ndr2* injected cells and adjacent cells involuted from periphery inward. From 6 hpf to 10 hpf, an obvious protrusion resembling blastopore lip (Xu *et al*., 2014) formed at the Nodal injected end, then Nodal explants gradually elongated, due to embryonic convergence and extension movements (Supplementary Movie 1). This elongation process is dose-dependently induced by Nodal. Note that 10 pg *ndr2* mRNA was sufficient to induce obvious elongation (Supplementary Fig. S1A). Therefore, except as otherwise mentioned, we used 10 pg of *ndr2* mRNA for subsequent experiments. Later, Nodal explants stopped elongating and curled before 24 hpf.

To characterize the cell types present in uninjected (control) and Nodal explants, we performed whole-mount *in situ* hybridization (WISH) for markers of the three germ layers (*sox2* and *otx2a* for the ectoderm, *tbxta* for the mesoderm and *sox32* for the endoderm) and we analyzed the impact of Nodal signaling on the expression of genes coding for signaling molecules known to be essential to early embryonic development. (Fig. 1B and Supplementary Figs. S1B, C). In agreement with previous reports (Xu *et al*., 2014), no transcripts of *fgf8a, wnt8a, ndr2* or *wnt11f2* are detected in control explants, while expression of these genes is observed in Nodal explants. In control explants, while the neuroectoderm marker *sox2* is expressed ubiquitously as early as 4 hpf there is no expression for mesoderm (*tbxta*) or endoderm (*sox32*) markers from 4 hpf to 10 hpf. This observation is in agreement with the anterior neural ectoderm being the default state of naive embryonic cells (Thisse *et al*., 2000) and the absence of expression of mesoderm and endoderm markers demonstrates the absence of germ layer inductive signal in the animal pole explants isolated at 512-cell stage. At 4 hpf, 1.25 hours after explants are taken, expression of *sox2* in Nodal explants is similar to the controls but is progressively restricted to the opposite end of the Nodal induced protrusion during the following 6 hours.

Expression of the mesoderm marker *tbxta* initiated early at 4 hpf in the Nodal injected site, then converged and extended in the central region of the explant over time. The endoderm marker *sox32* is detectable at 6 hpf, with patchy expression near the Nodal injected site, the endoderm involutes and migrates inside the explant before ending up at the opposite side at 10 hpf.

Collectively, these results suggest that zebrafish animal pole explants are a naïve and competent system, which is suitable for *ex vivo* study. By overexpressing *ndr2* mRNA at the animal pole, we artificially constructed a Nodal gradient and induced mesendoderm specification, as well as the patterning of neural ectoderm. Furthermore, graded Nodal expression induces morphogenesis and elongation in the explants, which is time-matched to that in the embryo.

### Nodal response genes and their expression dynamics during the onset of gastrulation

To achieve its function of regulating cell fates and morphogenesis, Nodal needs to activate groups of genes ranging from transcription factors to cytoskeletal components (Bennett *et al*., 2007; Coda *et al*., 2017; Dubrulle *et al*., 2015; Liu *et al*., 2011; Nelson *et al*., 2014). Both concentration and time of exposure play important roles in morphogen interpretation (Ashe & Briscoe, 2006; Dubrulle *et al*., 2015; Müller *et al*., 2012; Sako *et al*., 2016; Thisse *et al*., 2000).

To characterize the effect of Nodal concentration and duration on its downstream gene induction, explants injected with 2 pg, 6 pg, and 10 pg of *ndr2* mRNA were collected for performing bulk RNA-seq at 4 hpf, 5 hpf and 6 hpf (Fig. 2A). Over 4000 genes were up-regulated in Nodal explants compared to uninjected explants. These genes can be grouped into two distinct classes according to their expressing dynamics, with one group increasing and the other one decreasing over time (Supplementary Fig. S2A).

**Figure 2.**
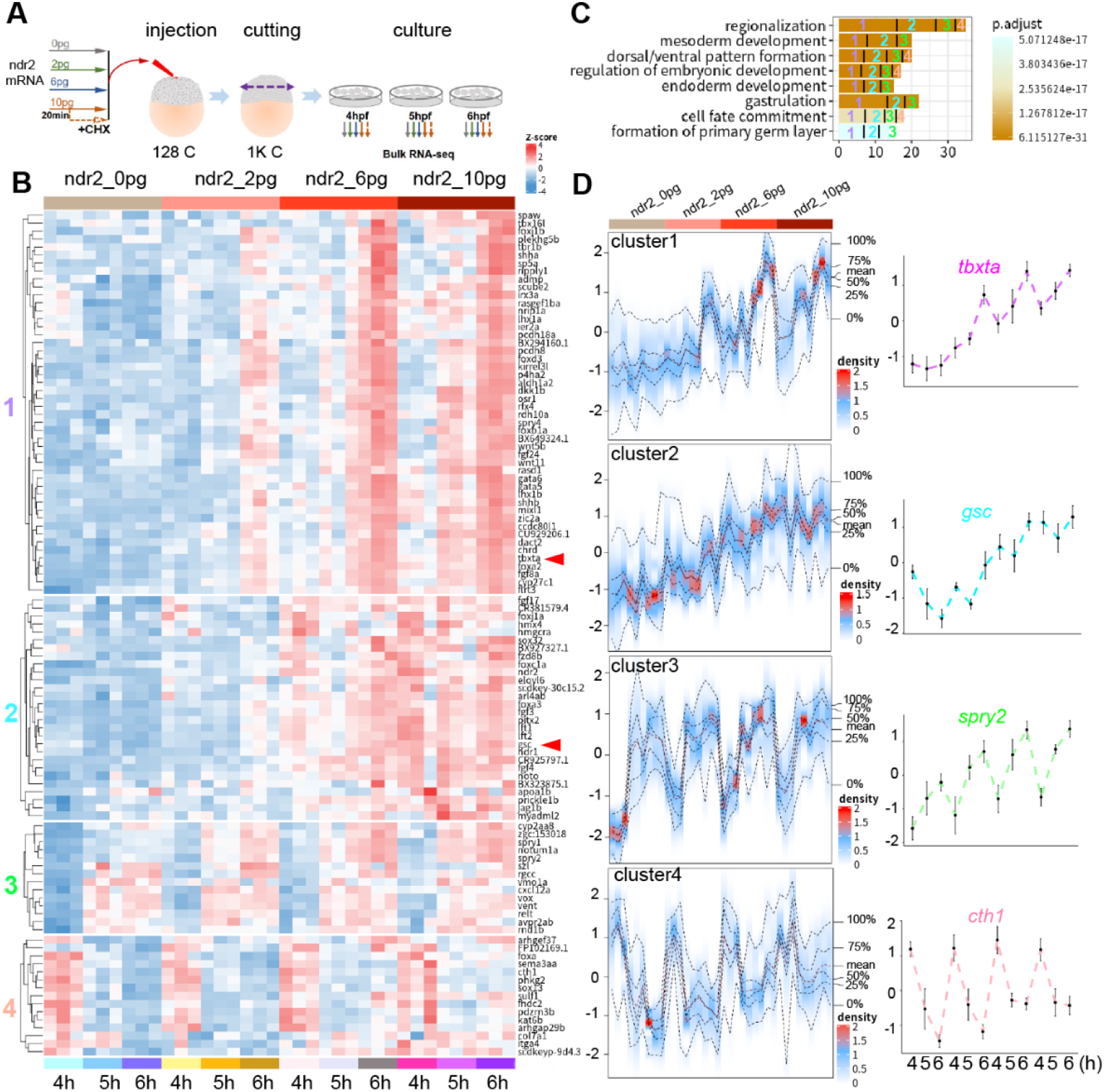
Identification of Nodal downstream genes and analysis of their expression dynamics. **A,** Cartoon illustrating experimental workflow of bulk RNA-seq in explants. 0 pg, 2 pg, 6 pg or 10 pg *ndr2* mRNA injected explants and 10 pg *ndr2* mRNA injected plus Cycloheximide (CHX) treated (dotted lines) explants were collected at 4 hpf, 5 hpf and 6 hpf for bulk RNA-seq. **B,** Gene expression heatmap of 105 Nodal immediate target genes aligned with samples of ascending Nodal concentration (according to **A** and labeled with colored bar at top of the heatmap, except CHX treated samples). Expression of each gene was z-scored across all genes in each sample. Target genes were clustered into 4 groups (indicated by numbers in the left) using k-means clustering. Red arrows showed the expression of *tbxta* and *gsc*. **C,** Top 8 enriched GO terms of 105 Nodal targets. Bar color indicates the logarithmic transformed adjusted P value. Each bar was proportionally partitioned with numbers refers to gene clusters of (**B**). **D,** Visualization of the expression dynamics of the 4 gene clusters of (**B**).

Nodal functions through a series of cascading reactions of its downstream genes (Schier, 2003). To identify Nodal immediate early genes (here called Nodal targets), we performed bulk RNA-seq for Nodal explants in the presence of the protein translation inhibitor Cycloheximide (CHX) (Fig. 2A). The efficiency of translation blocking was confirmed as CHX treatment could not influence the direct activation of *sox32* transcription by Nodal but blocked the expression of *sox17*, whose activation needs sox32 protein (Supplementary Fig. S2B). We first determined the differentially expressed genes between Nodal explants treated with CHX and uninjected explants. To exclude the artificial effects of CHX treatment, we overlapped those genes to the up-regulated genes in Nodal explants (without CHX, see details in **Methods**) compared to uninjected explants (Supplementary Figs. S2C-N). By this way, we identified 105 Nodal targets from three time points (Supplementary Fig. S2O). Among them, more than 60% have been reported as Nodal direct target genes in previous works [Supplementary Data 1 (Bennett *et al*., 2007; Liu *et al*., 2011; Nelson *et al*., 2014)]. These 105 Nodal targets can be clustered into 4 groups by K-means clustering (Figs. 2B, D). Among the 4 groups, genes of Cluster 1 and 2 were activated only when Nodal is present, suggesting that in the explants, Nodal is the only inducer of those two clusters. Genes from Cluster 3 and 4 were activated with or without Nodal, although the expression level of both clusters can be up-regulated by Nodal, suggesting that those genes are under the regulation of factors in addition to Nodal.

Interestingly, genes of Cluster 1 showed a clear pattern of increasing expression levels over time. The levels of gene expression induced by low concentrations for long periods and high concentrations for short periods of time by Nodal were similar. The characteristics of this cluster are consistent with a previous reviewed model of morphogen interpretation (Ashe & Briscoe, 2006). The expression level of Cluster 2 genes continuously increases with increasing Nodal, but didn’t show obvious changes over time. In previous work, *tbxta* was identified as a typical long-range Nodal target, which can be induced at low levels of Nodal and continuously increased over time (Dubrulle *et al*., 2015). By contrast, *gsc*, a short-range Nodal target, requires higher Nodal concentration to be activated (Dubrulle *et al*., 2015). *tbxta* and *gsc* were grouped into Cluster 1 and Cluster 2 respectively which fits well with previous study.

We constructed a protein-protein interaction network for Nodal targets (see **Methods**), and found that genes from Cluster 1 and Cluster 2 were the hub genes in the network (Supplementary Fig. S2P). GO analysis showed that the functions of these genes are mainly germ layer formation, dorsal-ventral patterning and regionalization, which is consistent with previous studies (Fig. 2C) (Feldman *et al*., 1998; Liu *et al*., 2018; Sampath *et al*., 1998; Shen, 2007). Cluster 1 and 2 contribute the most to all the GO categories. These results suggest that the genes from Cluster 1 and Cluster 2 are more important to the interpretation of Nodal signal and their Nodal-responsive elements can be used to study the temporal and spatial mechanisms of Nodal morphogen interpretation.

### Construction of a single cell response landscape of Nodal

Advances in single-cell RNA sequencing technology may allow understanding the cell types induced by Nodal at a higher resolution, and the origin and development of those cell types, as well as the interactions between different cell types. Towards this goal, 6 stages of Nodal explants were collected for single cell RNA sequencing. (Fig. 3A).

**Figure 3.**
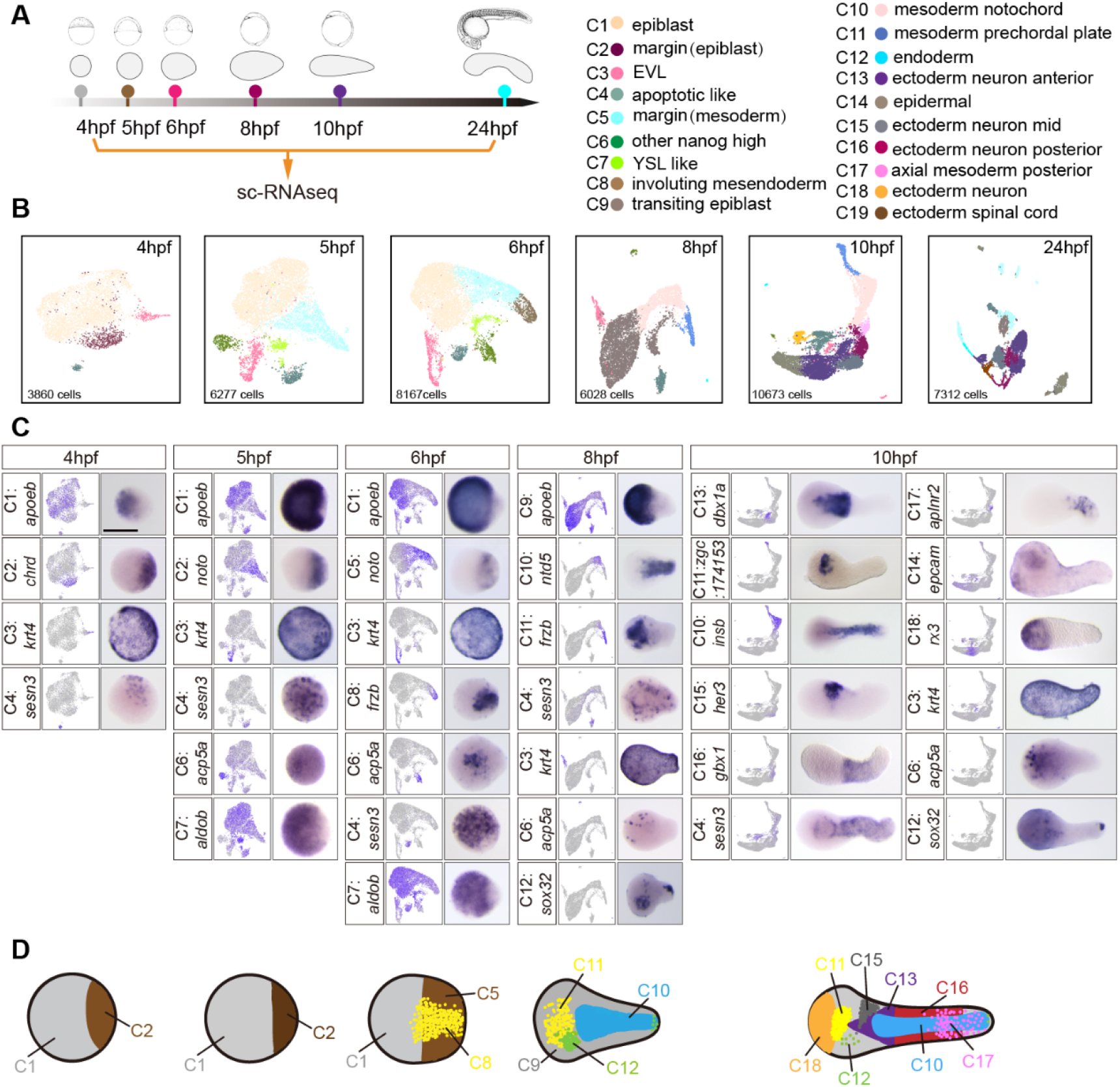
Single cell transcriptome analysis of Nodal injected explants. **A,** Schematic representation of experimental workflow. 10 pg of *ndr2* mRNA injected explants of 6 timepoints (4 hpf, 5 hpf, 6 hpf, 8 hpf, 10 hpf and 24 hpf) were collected for single cell RNA-seq. Representative explants of each timepoint are aligned chronologically in the second row, the corresponding embryos at each timepoint are also showed in the top row. **B,** UMAP plots for Nodal injected explants at each timepoint. Cells are colored according to their cell-type annotation inferred from expressed marker genes. **C,** The expression of selected marker genes from each cluster of Nodal injected explants were displayed by UMAP and WISH (from 4 hpf to 10 hpf).**D,** Cartoon plots show the expression pattern of main cell clusters in Nodal injected explants at indicated timepoints. Scale bar: 200 μm (**C**).

A total of 42,317 single-cell transcriptomes were collected after stringent quality control measurements, and visualized using a uniform manifold approximation and projection (UMAP) plot (Becht & McInnes, 2018) to show how cells cluster base on transcriptional similarity. 19 cell types from 6 developmental stages were identified and annotated by expression of marker genes (Supplementary Data 2). Similar to endogenous embryonic development, cell types in Nodal explants exhibited progressively increasing difference over time (Fig. 3B). To validate the cell types identified by scRNA-seq and investigate their properties and locations compared to wild-type embryo, we performed WISH for selected marker genes of each cluster at timepoints corresponding to the scRNA-seq in Nodal explants, uninjected explants and embryos (Fig. 3C and Supplementary Figs. S4A-E). We found that some cell types (C1: epiblast, C3: EVL, C4: apoptotic like, C6: other nanog high, C7: YSL like, C9: transiting epiblast, C14: epidermal and C15: ectoderm neuron mid) could be observed both in Nodal explants and in uninjected explants, suggesting that those cell types appear spontaneously in a Nodal independent manner. The other cell types were all exclusively found in Nodal explants. Among the spontaneous cell types, epiblast (C1, C9) and neural ectoderm (C15), which was ubiquitously distributed in the entire uninjected explants, were progressively restricted to a specific territory in Nodal explants, indicating that those cell types could be patterned even if they are not directly induced by Nodal. EVL and epidermal cells were largely unaffected by Nodal. Interestingly, we found a group of cells (C4) expressing genes regulating apoptosis emerged as early as 4 hpf in Nodal injected and uninjected explant and wild-type embryo, which were also found in previous studies (Farrell & Wang, 2018; Satija *et al*., 2015; Wagner *et al*., 2018). We also found two groups of cells (C6 and C7) that, although they are not clustered together, both express YSL genes. We term these clusters “YSL like” (C7) and “other nanog high” (C6), which corresponds to a previously found cluster (Wagner *et al*., 2018). As early as 4 hpf, *chordin*, indicating the dorsal margin/organizer (C2), can be detected in Nodal explants. Starting from 5 hpf, the mesoderm (C5) is specified at the Nodal injected site, then the separation of involuting mesendoderm (C8) (Figs. 1B and 3B, C) and margin mesoderm (C5) become apparent by both the UMAP plot and WISH.

From mid to late gastrulation (8 hpf, 10 hpf), prechordal mesoderm (C11) and notochord mesoderm (C10) became transcriptionally and positionally distinct in Nodal explants. From the onset to the end of gastrulation (6 hpf, 8 hpf, 10 hpf), the endoderm involuted and migrated to the anterior (the end opposite the site of Nodal injection) of the Nodal explants. Interestingly, some of the *sox32* positive cells do not involute and stay at the tip of the posterior end (the Nodal injection site), which we speculate might resemble the dorsal forerunner cells (DFCs). At the end of gastrulation, besides prechordal and notochordal mesoderm, a group of posterior axial mesoderm cells (tailbud, C17) can be detected at the end of Nodal explants. Intriguingly, at this stage, the anterior (C18, C13), middle (C15) and posterior (C16) neural ectoderm were sequentially ordered along the anterior-posterior axis of Nodal explants, while in the uninjected explant we didn’t observe any of those cell types except neural ectoderm mid (C15). Overall, the dynamics and location of the main cell types in Nodal explants (as summarized in the cartoon diagram, Fig. 3D), together with lineage analysis (Supplementary Figs. S3A, B) show that Nodal explants could, to some extent, reflect the spatiotemporal characteristics of embryonic development.

To fully investigate what tissues and organs can be initiated and organized by Nodal alone, we combined scRNA-seq data from Nodal explants and embryos (Wagner *et al*., 2018) at 24 hpf (see **Methods**). In total 33 cell clusters were identified after unsupervised clustering (Stuart *et al*., 2019) of the combined dataset (Supplementary Fig. S3C and Supplementary Data 3). All clusters were annotated based on previously published information (Supplementary Data 3). We found that cells from Nodal explants can be co-clustered with embryonic cells (Supplementary Figs. S3D, F). However, some clusters were nearly absent in Nodal explants, including cluster 17 (finbud; heart), cluster 24 (macrophage; leukocyte), cluster 30 (differentiating neurons), cluster 31 (roofplate), cluster 23 (ionocyte), cluster 29 (rohon beard neuron), cluster 6 and cluster 20 (erythroid cell). While for cluster 13 (erythroid cell), cluster 28 (hatching gland) and cluster 15 (mesoderm; PSM), the proportion of cells from Nodal explants was much higher than that from embryos. Surprisingly, although cluster 13 was annotated as erythroid cells in embryonic dataset, cells from Nodal explants in cluster 13 did not express any erythroid cell genes, but rather highly expressed genes of muscle cells (Supplementary Fig. S3E). A previous study reported that a subset of hematopoietic stem cells are derived from the somites in zebrafish (Qiu *et al*., 2016), thus we speculate that the Nodal explants cells in cluster 13 could be this subset of hematopoietic stem cells, which cannot further differentiated into mature erythroid cells due to lack of some essential signals.

Expression of selected marker genes from 33 cell clusters and some organ primordia (heart: *hand2;* gut: *gata5* and *gata6*) were checked in both Nodal explants and embryos at 24 hpf (Supplementary Figs. S3C and S4G-J). Surprisingly, we didn’t find any internal organs derived from endoderm. Although the tSNE map showed that there are some pharyngeal pouch cells in the Nodal explants, we didn’t observe any *pax1b* expression by WISH, possibly because there are too few of these cells. For tissues derived from mesoderm, only hatching gland and tail bud were found in Nodal explants, but there is no tissue of the trunk region such as the heart and blood vessels. Remarkably, the anterior part of Nodal explants contains mainly neural ectoderm cells, including forebrain, midbrain, hindbrain, eye and spinal cord, and these cell types are properly organized along the anterior-posterior axis and highly similar to the pattern of embryonic head.

### Reconstruction of pseudospatial axes of Nodal explant

One of the advantages of explants over embryos is that it is a simplified model, morphologically the Nodal explant has only two axes: one along the anterior-posterior and one radially from this axis. To better understand the relationships between the pattern of cell types and position of gene expression under Nodal regulation, we constructed two pseudospatial maps from our single cell RNA-seq data of Nodal explants at before (4 hpf and 5 hpf) and the onset of gastrulation stages (6 hpf).

Firstly, for the anterior-posterior axis, we performed dimensionality reduction to the embeddings of Nodal response cells (C1: epiblast, C5: margin mesoderm, C8: involuting mesendoderm) in the 3-D UMAP plot of Nodal explants (Fig. 4A) by LDA (Linear Discriminant Analysis) algorithm, thus constructing a linearized two-dimensional plot with orderly arranged mesendoderm, mesoderm and ectoderm which we call a “pseudospatial axis” (Fig. 4B). Then we looked at the distribution of the expressions of the previously identified 105 Nodal targets along this pseudospatial axis at 4 hpf, 5 hpf and 6 hpf (Supplementary Figs. S5A, B and Fig. 4B). We found that the expression cascade of these genes along the pseudospatial axis gradually showed a gradient pattern, in which two expression thresholds could be observed at 6 hpf. Note that these two gene expression thresholds align nicely with the boundary of different cell types, indicating that the gene expression thresholds formed by morphogen-initiated gene regulatory networks, further clarified the boundaries between different cell types.

**Figure 4.**
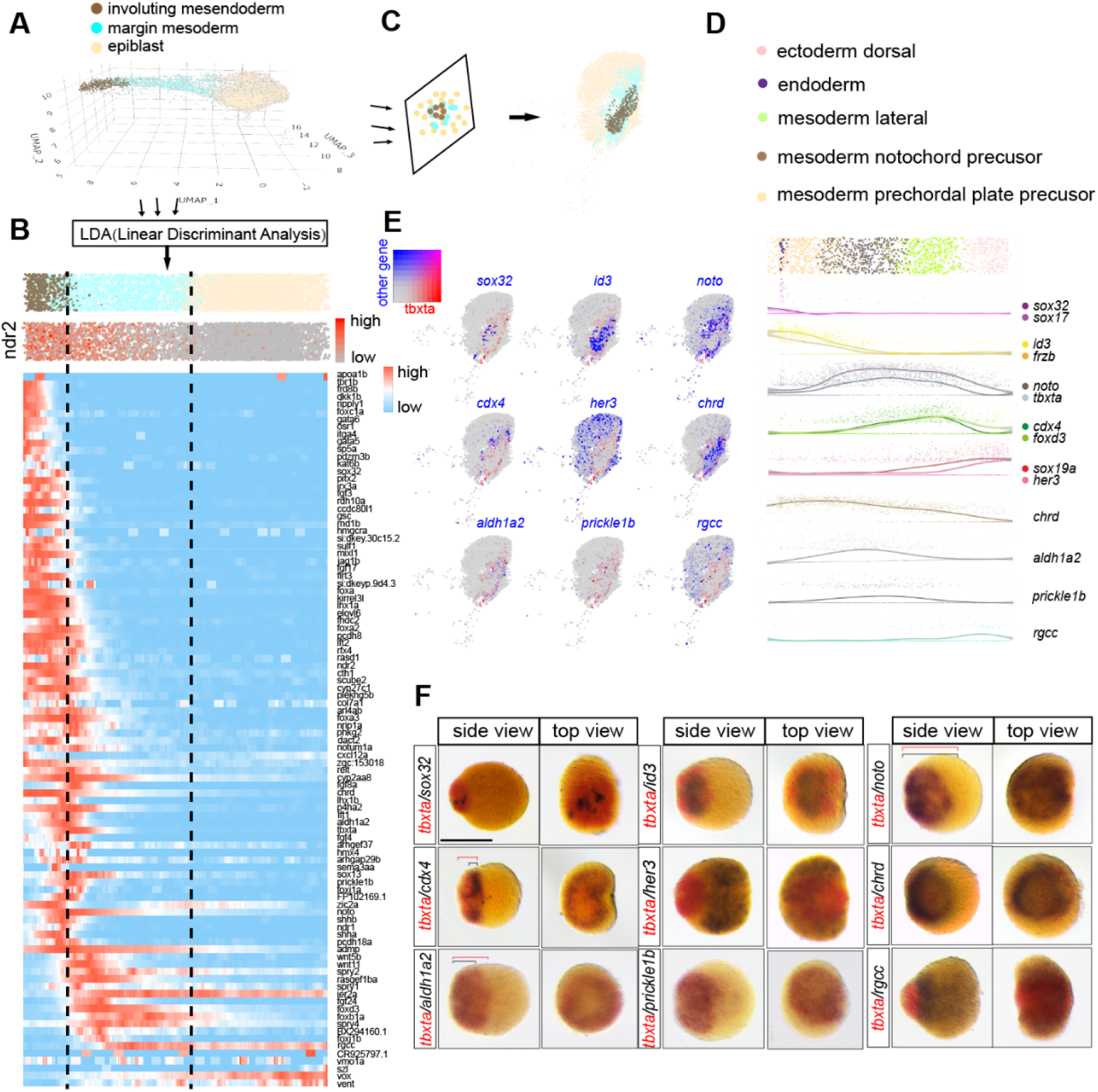
Construction of pseudo-spatial axis of Nodal injected explant at gastrulation stage. **A,** 3-D UMAP representation of three germ layer cells in Nodal injected explants at 6 hpf. **B,** A linearized two-dimension plot was reconstructed from the 3-D UMAP plot in (**A**) using linear discriminant analysis (LDA, see Methods for details), which we term it a pseudo-spatial axis. Graded expression of *ndr2* mRNA are observed in this axis. Heatmap shows the expression cascades of Nodal targets along this axis. **C,** Cells were projected onto a UMAP_2 versus UMAP_3 plane, for constructing a flatted 2-D UMAP plot. **D,** Pseudo-spatial axis of *ndr2* positive cells in 6 hpf Nodal injected explant. The *ndr2* positive cells were taken out and annotated with subdivided cell clusters. The expression profiles of cluster markers (endoderm: *sox32, sox17*; prechordal plate: *id3, frzb*; notochord: *noto, tbxta*; lateral mesoderm: *cdx4, foxd3*; ectoderm: *sox19a, her3*) and selected Nodal targets (*chrd, aldh1a2, prickle1b* and *rgcc*) are shown along the pseudo-spatial axis. **E,** Expression profiles of *tbxta* (red) and selected genes (blue) on the flatted UMAP plot from (**C**). **F,** Double color WISH shown the expression pattern of *tbxta* (red) and genes (blue) from (**E**) in Nodal injected explant at 6 hpf. Scale bar: 200 μm (**F**).

We also noticed graded expression of *ndr2* along this pseudospatial axis (Fig. 4B), which may reflect the Nodal activity gradient to some extent (van Boxtel *et al*., 2015). Since *ndr2* positive cells received the highest level of Nodal, and thus were the most affected by Nodal signal, we picked out these cells and constructed a cell atlas of 5 stages. Cell types showing rare cell populations such as endoderm cells now can be revealed as a separated cluster in 6 hpf Nodal explants (Supplementary Fig. S5C). We also constructed a pseudospatial axis of *ndr2* positive cells from 6 hpf Nodal explants with endoderm, mesoderm prechordal plate precursor, mesoderm notochord precursor, mesoderm lateral and ectoderm dorsal, which were orderly arranged along the pseudospatial axis consistent with the endogenous spatial distribution of these cell types. The expression positions of marker genes from each cluster were also aligned with their corresponding cell types along this pseudospatial axis (Fig. 4D).

For the axis that extends radially from the center, we flattened the 3-D UMAP plot into a “disk shaped” 2-D UMAP plot, in which the mesendoderm, mesoderm and ectoderm are arranged from the center outward. For this application, we tried to visualize the expression patterns of marker genes of different cell types and several Nodal targets in these two coordinate systems, and these *in silico* patterns were further validated by WISH (Figs. 4D-F). In summary, we have constructed a molecular pseudospatial axis of Nodal explants that resembles the animal-vegetal (An-Vg) axis of zebrafish embryos at the onset of gastrulation. The molecular features of the pseudospatial axis could help us to better understand how a morphogen regulates cell specification in a concentration dependent manner.

### Different Nodal downstream targets can induce different portions of the body axis

Nodal signaling is essential for the formation of the dorsal organizer (Feldman *et al*., 1998; Gritsman *et al*., 2000), and overexpression of Nodal at the ventral side of early embryo can induce a secondary axis [Fig. 6B and (Lustig *et al*., 1996)], showing that Nodal itself plays an organizer function. This function is achieved by its downstream genes, such as Goosecoid (Gsc) (Toyama *et al*., 1995), which was identified as the first organizer-specific gene (Cho *et al*., 1991) and can elicit complete secondary axes when expressed ventrally in many animal models including zebrafish (Dixon Fox & Bruce, 2009). Besides Gsc, several other genes like Chordin and Dkk1 have also been shown to have axis induction ability (Dixon Fox & Bruce, 2009; Kazanskaya *et al*., 2000), and both *chordin* and *dkk1b* were identified as Nodal targets in this study.

To fully understand the molecular mechanism of the organizer function of Nodal, we isolated *ndr2* positive cells from Nodal explants (4 hpf to 10 hpf) for enrichment of the cell types under high Nodal signaling, which includes endoderm and axial mesoderm such as prechordal plate (Supplementary Fig. S5C and Supplementary Data 4). We then reconstructed developmental trajectories of *ndr2* positive cells from 4 hpf to 10 hpf using the URD algorithm (Farrell & Wang, 2018). The cell state tree followed the specification of 8 final cell clusters including: neural plate posterior (NPP), neural plate anterior (NPA), neuromesodermal progenitor (NMP), paraxial mesoderm (PM), notochord (NT), tailbud (TB), prechordal plate (PP) and endoderm (Endo) (Figs. 5A, B).

**Figure 5.**
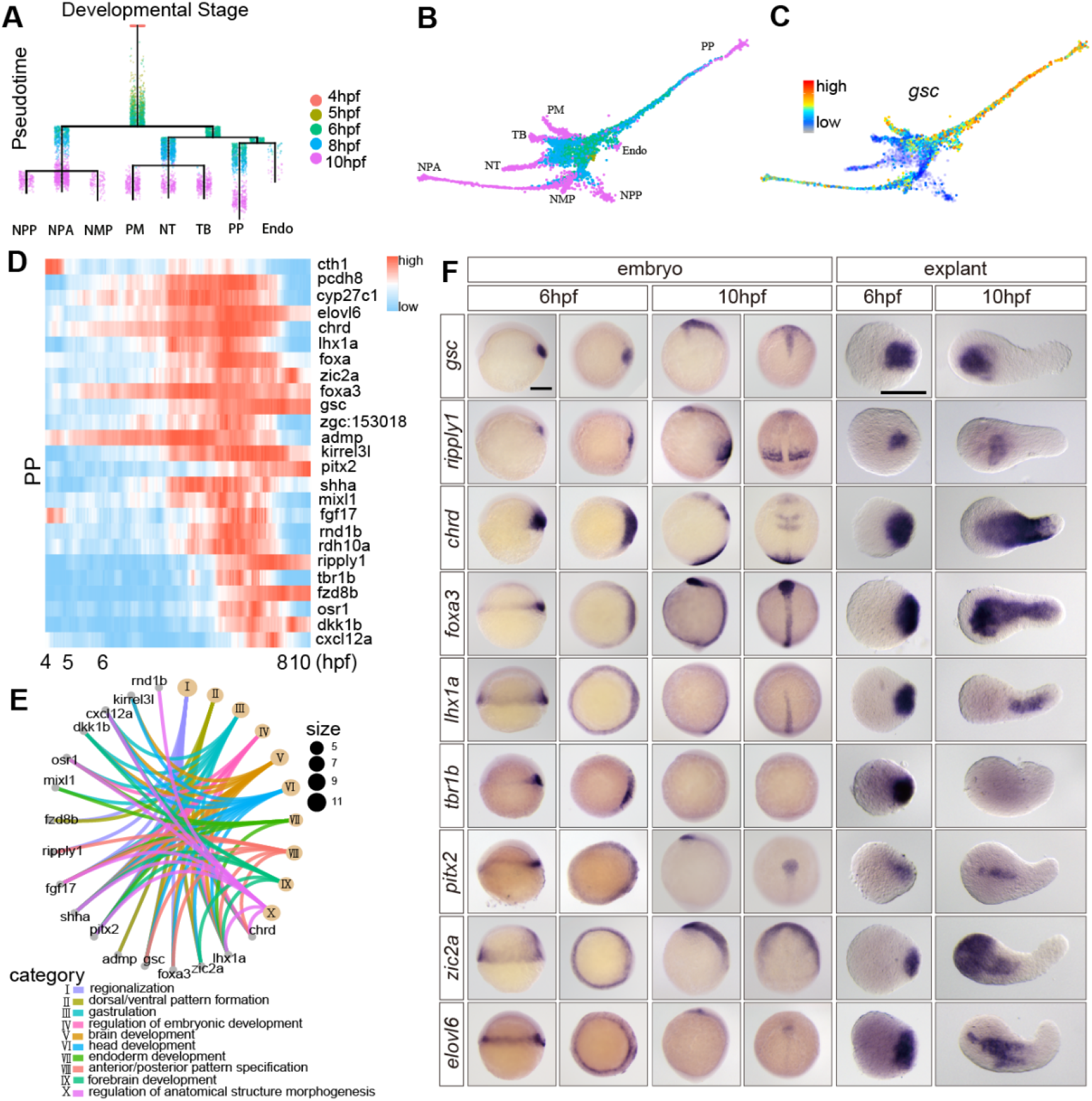
Expression features of the 25 Nodal targets which highly expressed in PP lineage. **A,** Dendrogram layout of URD differentiation tree of *ndr2* positive cells. Cells are colored by developmental stages. (NPP, neural plate posterior; NPA, neural plate anterior; NMP, neural mesoderm progenitor; PM, paraxial mesoderm; NT, notochord; TB, tailbud; PP, prechordal plate; Endo, endoderm). **B,** Force-directed layout of URD differentiation tree, optimized for 2-D visualization. Cells are colored by developmental stages (same to **A**). **C,** Expression profiles of *gsc* on the differentiation tree, *gsc* is highly expressed in PP. **D,** Pseudotemporal gene-expression cascades of PP cell lineage. Lineage specific Nodal targets were recovered by differential expression testing from URD and used for construction of the expression cascades, 25 Nodal targets are identified. **E,** cnetplot (see **Methods**) showing the linkages of PP lineage Nodal targets and enriched top10 GO terms. **F,** In situ hybridization of the selected genes from 25 Nodal targets in Nodal injected explants and embryos at 6 hpf and 10 hpf. Scale bar: 200 μm (**F**).

To identify potential organizer function genes downstream of Nodal, we used *gsc* as a reference and explored the expression dynamics of Nodal targets on the cell state tree (Supplementary Figs. S5D, E). Consistent with previous studies (Farrell & Wang, 2018; Schulte-Merker *et al*., 1994), we found *gsc* highly expressed in the PP (Fig. 5C) along with 24 Nodal targets (Fig. 5D) whose GO terms are enriched for function in tissue specification and patterning (Fig. 5E). We confirmed the expression patterns of all these 25 Nodal targets by WISH at 6 hpf and 10 hpf (Fig. 5F and Supplementary Fig. S6A), both in Nodal injected explants and in wild type embryos. To investigate the axis induction capabilities of these genes, we injected their mRNAs separately (with fluorescent dye) into one blastomere of zebrafish embryos at 16-32 cell stage. The embryos with ventrally localized fluorescent clones at 6 hpf were selected (Fig. 6A) for induction of a secondary axis at 12 hpf. We found that among the 25 genes, 14 of them can induce a complete or incomplete secondary body axis, as evidence by the morphology of live embryos and WISH of *six3b, egr2b, shha, pax2a* and *cdx4* (Figs. 6B, C and Supplementary Figs. S6B, C). Of these, only *gsc* can induce a complete axis. *ripply1* showed strong induction capabilities but without notochord. Genes such as *chrd, foxa3, lhx1a, tbr1b, pitx2, foxa, elovl6* can induce different portions of the axis (Figs. 6B, C and 7A). 11 other Nodal targets did not show any axis induction capabilities (Supplementary Fig. S6D). We ranked the induction capabilities of these genes by analyzing the induced tissues and the length of the induced axes (Fig. 7A).

**Figure 6.**
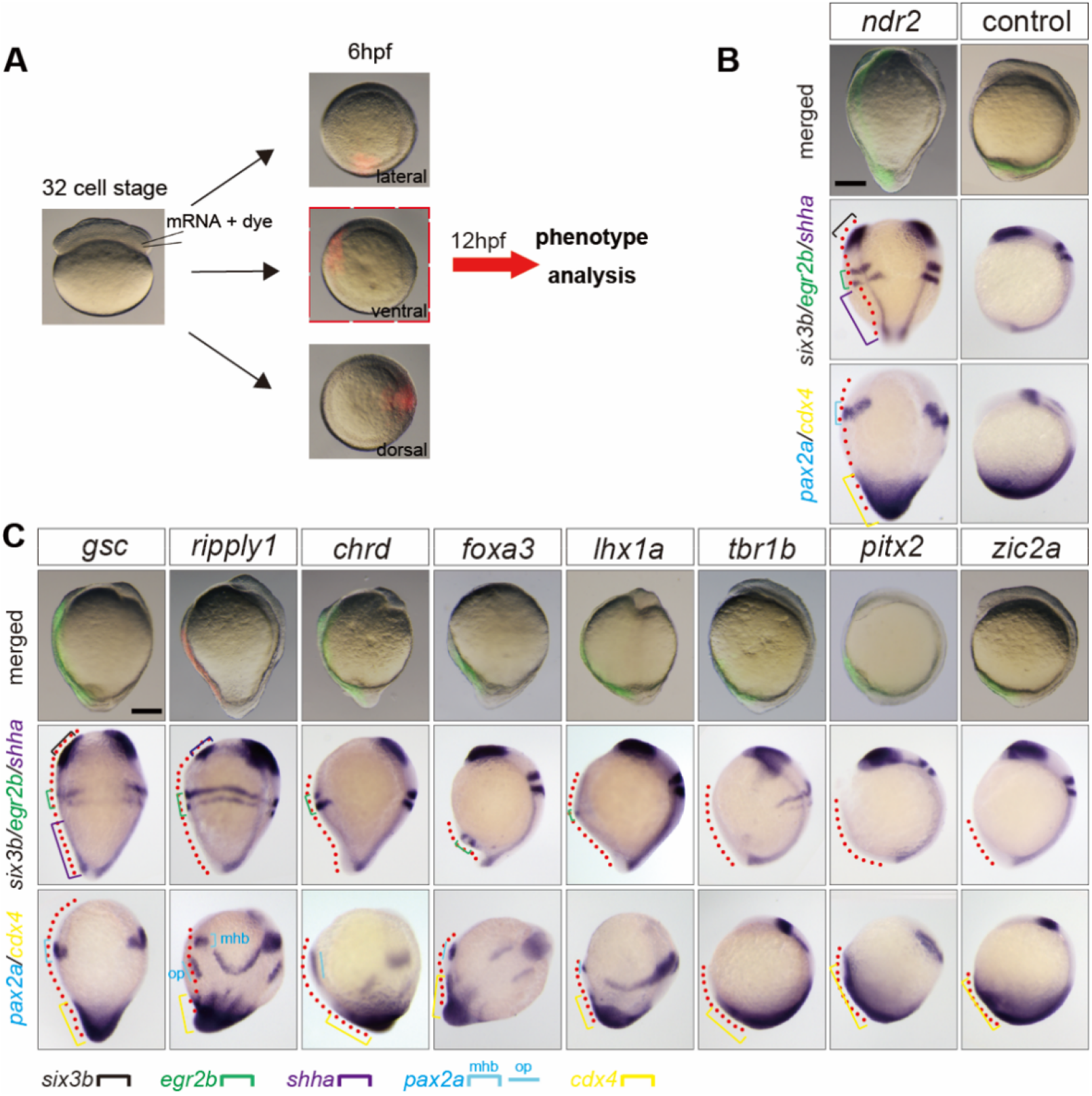
Different Nodal targets can induce complete or partial secondary axes when ventrally overexpressed. **A,** Schematic diagrams show the strategy of ventral overexpression. **B,** Upper: merged images of live zebrafish embryo injected with *ndr2* (5 pg) and dye (control). Fluorescein indicates the clones of the injected blastomere at 32 cell stage. Embryos injected with dye do not show secondary axis structures at 12 hpf. Middle: In situ hybridization of *six3b* (forebrain), *egr2b* (hindbrain), and *shha* (notochord) show the induced secondary axis structures at 12 hpf. Bottom: In situ hybridization of *pax2a* [midbrain hindbrain boundary (mhb) and otic placode (op)] and *cdx4* (posterior mesoderm and posterior neural ectoderm) show the induced secondary axis structures at 12 hpf. Red dash lines indicate induced secondary axes. **C,** Upper: merged images of live zebrafish embryo injected with indicated mRNAs. Middle: In situ hybridization of *six3b, egr2b* and *shha* at 12 hpf. Bottom: In situ hybridization of *pax2a* and *cdx4* at 12 hpf. Scale bar: 200 μm (**B** and **C**).

**Figure 7.**
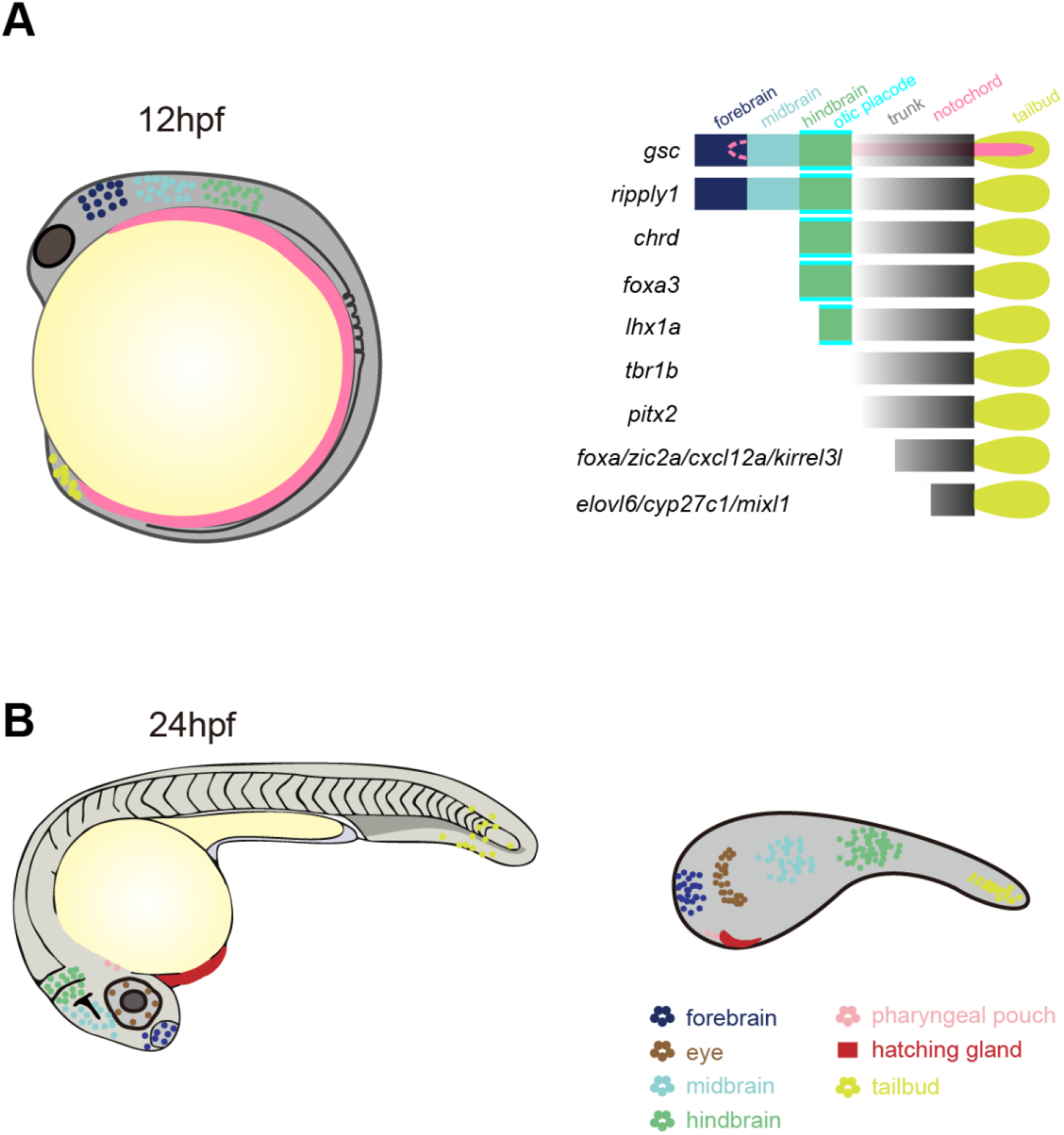
Model of Nodal function during zebrafish early development. **A,** Cartoon illustrating the different induction capabilities of the 14 Nodal targets. *gsc* showed strongest induction capabilities which can induce a complete secondary axis including head, trunk and notochord. *ripply1* can induce most structures except notochord. Both *chrd* and *foxa3* can induce part of head structure (hindbrain and otic placode) and trunk. *lhx1a* can induce part of hindbrain, otic placode and trunk. *tbr1b, pitx2, foxa, zic2a, cxcl12a, kirrel3l, elovl6, cyp27c1* and *mixl1* can induce trunk with different length. **B,** Cartoon illustrating the major cell types identified in Nodal injected explants and their corresponding positions in zebrafish embryos at 24 hpf. Anterior neuroectoderm is well-patterned in Nodal injected explants including forebrain, midbrain and hindbrain. Lateral mesoderm is absent, while the axial mesoderm such as hatching gland and tailbud are found located in anterior and posterior end of the explant respectively. Except for the anterior endoderm (pharyngeal pouch), other endoderm cell types are hardly detected in Nodal injected explants.

### Ripply1 and Gsc regulate separation of the Mesoderm and Endoderm

Previous work has shown that endoderm and axial mesoderm are both induced by high levels of Nodal signal (Barone *et al*., 2017), but the molecular mechanisms for separation of these two lineages is not yet fully understood. On the cell state trajectory tree constructed from the scRNA-seq data of our Nodal explants, the process of endoderm and prechordal plate bifurcation from a common precursor is clearly observed (Fig. 5A). We performed differential analysis for endoderm and PP cells and found, several Nodal targets with differential expression (Fig. 5D and Supplementary Figs. S7B, C). We constructed a branchpoint plot separating the PP and endoderm (Supplementary Fig. S7D). Consistent with previous results, PP markers (*gsc, ripply1*) and endoderm markers (*sox32, sox17*) were highly expressed in the two different cell types (Supplementary Fig. S7E). It has been shown that endoderm differentiation could be suppressed by upregulation of *gsc* (Sako *et al*., 2016). As Ripply1 and Gsc are both transcriptional repressors (Kawamura *et al*., 2005; Sako *et al*., 2016) and have a similar function in induction of a secondary axis. We hypothesized that *ripply1* may also suppress endoderm. To test this hypothesis, we injected *ripply1* mRNA at the 1-cell stage and found that endoderm induction decreased as shown by WISH of *sox32* and EGFP expression driven by sox17 promoter. To specifically overexpress *ripply1* in the endoderm, we constructed a plasmid that contains *ripply1* cDNA downstream of the sox17 promoter. Injection of this plasmid at 1 cell stage also resulted in a decrease in endoderm (Supplementary Figs. S7F, G). Taken together, these results demonstrated that Ripply1 may act as an endoderm repressor similar to Gsc.

## Discussion

In this study, we used the animal pole explant of zebrafish blastula as a research model for morphogen mediated patterning. We firstly confirmed that the animal half of the zebrafish animal cap is a relatively naive system which contains very limited signals. In the absence of external signal, explants will not undergo obvious morphogenesis, nor will cell types form besides default anterior neural ectoderm. This suggests that a group of homogeneous cells does not have the intrinsic potential to break symmetry and will not spontaneously undergo morphogenesis through self-organization. However, this explant can respond well to exogenous signals such as Nodal. In the presence of a Nodal gradient, we found that explants undergo a series of cell movements which is very similar to embryonic gastrulation: firstly, a group of cells located in the site of high Nodal activity collectively involutes into the interior of the explant and forms endoderm and prechordal plate; then an obvious convergence and extension movement of the entire explant makes it significantly elongate, and results to the formation of a notochord in the middle.

In recent years “embryoid” and “gastruloid” models have received great attention as new research models (Beccari *et al*., 2018; Moris *et al*., 2020; Rossi *et al*., 2020; Veenvliet *et al*., 2020), our work shows that a single Nodal gradient can robustly induce a “gastruloid” with different germ layers and anterior-posterior pattern from a group of naïve cells of zebrafish, and provides good guidance for using a morphogen gradient to induce “embryoids” or “gastruloids” from embryonic stem cells of mouse or human.

Nodal is a TGF-β-superfamily member that binds to its receptors and causes the phosphorylation of smad2/3 (Hill, 2018). The consensus is that Nodal as a morphogen can form a gradient that results in a corresponding gradient of pSmad2/3, which then activates the expression of a series of downstream genes. The interaction of these genes further leads to the specification of different cell fates and cell movements. The animal pole explant does not contain any endogenous Nodal signal and is thus an ideal system for identifying the downstream genes induced by it. Using translational inhibition and bulk RNA-seq, we identified the first wave of 105 genes induced by Nodal at the onset of gastrulation. Interestingly, among those 105 genes, the expression dynamics of one group of genes showed high sensitivity to Nodal concentration and another group was more sensitive to Nodal exposure time. This is consistent with previous hypotheses that there are concentration dependent and exposure time dependent mechanisms in the interpretation of morphogens (Dubrulle *et al*., 2015). Future studies will be needed to determine how concentration and exposure duration are mechanistically distinguished.

Technical advances in transcriptomic analysis have greatly advanced developmental biology, especially in studying the continuous dynamic processes of cell lineages at higher resolution, identifying key genes that regulate cell fate decisions, and understanding the relationship between cell fate and its spatial distribution. Using the single-cell transcriptome data of Nodal explants from 6 developmental stages, we developed an algorithm to reduce the dimensions of these data along the long axis and the axis from the center to the periphery of the explant, thus we obtained two coordinate systems which we consider to be spatially similar to the Nodal induced patterning and can reflect the gradient of Nodal activity to a certain extent. The expression of Nodal target genes is distributed along these two coordinate systems in a graded manner, and this gradient is progressively established during development, which supports the hypothesis that the gene regulatory network regulates their expression thresholds and further determines the cell fates.

Through the construction of the single cell atlas and WISH validation, we found that Nodal can pattern explants by inducing expression of genes such as *chordin* as early as at 4 hpf, which is consistent with *in vivo* embryonic development. From 5 hpf, three germ layers are progressively induced and the patterning becomes more and more clear along the anterior-posterior axis of the explant. At 10 hpf, the endoderm and mesoderm cells that were originally at the posterior end migrate to the anterior end inside the explant through an involuting movement, at the same time, a head structure with anterior-posterior patterning is formed at the adjacent position on the periphery of the explant. Surprisingly, in the 24 hpf Nodal explant, the only endodermal tissue is the pharyngeal pouch, and mesoderm derived tissues are hatching gland and tail bud. Instead, the main population of the Nodal explant is a well anterior-posterior patterned head structure.

On the developmental trajectory tree of Nodal positive cells, 25 Nodal targets are highly enriched in prechordal plate including *gsc* which was the first identified organizer-specific gene. Among the 25 Nodal targets, 14 of them can induce a secondary axis with different tissue structures when they are ventrally overexpressed (Figs. 6, 7A and Supplementary Figs. S6B-D). At the same time, Ripply1 and Gsc which have the strongest ability to induce secondary axis (Fig. 7A) show redundant function in the suppression of endoderm specification (Supplementary Figs. S7F, G). Additionally, during the process of the separation of endoderm and prechordal plate (Supplementary Fig. S7D), we found that cadherin genes are differentially expressed in prechordal plate (*cdh1*) and endoderm (*cdh2* and *cdh6*) (Supplementary Fig. S7A). The differential cadherin expressions suggest that an adhesion code may be involved in the germ layer specification during development which support previous studies that cell-cell contact formation and cell-fate specification are highly related (Maître *et al*., 2012; Tsai *et al*., 2020).

In summary, this study systematically analyzed expression dynamics of Nodal targets, cell fate specification and patterning induced by Nodal in a naive system. We found that Nodal alone is sufficient to induce the dorsal mesoderm and anterior endoderm formation, and a well-organized head structure (Fig. 7B). Our work provides new understandings and a rich resource of Nodal induced targets and cell types, which will greatly help in identifying the mechanisms of morphogenesis such as involution and convergence and extension induced by it.

## Methods

### Animals

Zebrafish strains were conducted to standard procedures and experimental procedures were approved by the Institutional Review Board of Zhejiang University.

### Engineering of Nodal injected explants

Nodal injected explants were generated as follows: injecting 10 pg of nodal-related 2 (ndr2, cyclops) mRNA into one of animal pole blastomeres at the 128-cell stage, using a syringe needle (27G SUNGSHIM MEDICAL CO.,LTD) as a tool, the animal pole region of the embryos was cut from the blastula at the 1k-cell stage in a Petri dish coated with 1.5% agarose filled with Dulbecco’s Modified Eagle Medium: Nutrient Mixture F-12 (DMEM/F-12) Media with 10 mM Hepes, 1x MEM Non-Essential Amino Acids and supplemented with 7 mM CaCl2, 1 mM sodium pyruvate, 50μg/ml gentamycin, 2-mercaptoethanol 100μM,1x antibiotic-antimycotic (Gibco) and 10% Serum Replacement. Animal pole explants corresponding roughly to half of the blastula were allowed to form a sphere in this dish for a few minutes and then placed in another Petri dish containing fresh culture medium and incubated at 29 °C.

### Immunostaining and quantification of pSmad2 staining

Anti-pSmad2/3 (8828, Cell Signaling Technologies) antibody was used for Immunostaining experiments. Immunostaining for pSmad2/3 was carried out as described previously (van Boxtel *et al*., 2015). Confocal laser scanning microscopy was performed by high-resolution light sheet microscope (Light Innovation Technology Limited, LiTone LBS Light Sheet Microscope). The glass capillary was inserted into the microscope sample stage, then the plunger was gently pushed down such that the agarose-mounted-explants were just outside the capillary. Each explant was then targeted and orientated in EPI fluorescent mode with a 4x objective, and finally the 3-D stacks of each explant was captured in the light-sheet mode using a 25x water immersion objective (Nikon CFI75 25XW, N.A.=1.1) at 250nm step size with minimum laser excitations at 488 nm. 3-D distance between cells are measured using a customized imaging analysis pipeline for 3-D cell segmentation and quantification using the well-established Random Sample Consensus (RANSAC) method. The software was written in C++ and executed on a workstation with intel i7 6800k processor and 32 G RAM. The cell with highest P-Smad2 signal was assigned as central cell and it’s P-Smad2 intensity was normalized to 1. For other cells, the distance and P-Smad2 intensity were calculated by comparing with the central cell. The fitting curve was calculated and plotted by R (https://www.r-project.org).

### Sample preparation for bulk RNA-seq

10 pg, 6 pg, 2 pg of *ndr2* mRNA was injected to one blastomere at animal pole of zebrafish embryo at 128 cell stage. Uninjected samples were used as control. For the samples injected with10 pg *ndr2* mRNA, 20 min after the injection, half of the samples were treated with Cycloheximide (MCE, CAS No. 66-81-9) at 50 μg/μl until harvest. All embryos were incubated in 0.3x Danieau buffer until 1k stage, then, were transferred to Dulbecco’s Modified Eagle Medium. Animal pole blastomere corresponding roughly to half of the blastula were cut down and incubated to 4 hpf, 5 hpf, 6 hpf (corresponding to stage-matched control embryos) separately. All the samples (45 samples, three replicates for each treatment) were send to Novogene (Novogene Bioinformatics Technology Co., Ltd, Bejing, China) for the RNA extraction, cDNA library construction and sequencing. Paired-end sequencing (Novaseq6000, 150-bp reads) was performed.

### Sample preparation for single-cell RNA-seq

Nodal injected explants (injected with 10 pg *ndr2* mRNA) were harvested at 4 hpf, 5 hpf, 6 hpf, 8 hpf, 10 hpf, and 24 hpf (corresponding to stage-matched control embryos). 50-100 explants were transferred to 1.5 ml LoBind microcentrifuge tubes (Eppendorf 022431021) that had been rinsed with 10% BSA (Sangon Biotech Co., Ltd, A0903-5g)/Ringer’s solution (5 mM KCl, 0.12 M NaCl) for 20 min at room temperature. The supernatant was removed, and Trypsin-EDTA Solution (Beyotime, C0201, 400 μl for 24 hpf explants, 200 μl for other timepoint explants) was added to dissociate explants (30 min for 24 hpf explants, 20 min for other timepoint explants). Pipetting the entire volume of solution up and down several times through a P200 tip every 5 min. Add 200 μl 1% BSA/ Ringer’s solution to the tube after dissociating all the explants. The mixture was centrifuged to pellet cells (0.2 g, 4 °C, 5 min, Thermo, MICROCL 17R), then the supernatant was removed, and cells were resuspended in 200 μl 1% BSA/Ringer’s solution. Repeat upper procedures, and finally, the cells were resuspended in 200 μl 0.05% BSA/Ringer’s solution. Cell density was quantified manually using Hemacytometers (QIUJING Co., Ltd) and cell viability was analyzed using PI staining. Libraries were prepared using Chromium Controller and Chromium Single Cell 3’Library & Gel Bead Kit v2 (10x Genomics, PN-120237) according to the manufacturer’s protocol for 10000 cells recovery. Final libraries were sequenced on the Illumina Novaseq6000 (Genergy Biotechnology, Shanghai Co., Ltd).

### Processing and analyzing bulk RNA-seq data

Bulk RNA-seq datasets were aligned to the zebrafish GRCz11 genome using STAR (Dobin *et al*., 2013) and the reads mapped to multiple genomic location were removed. Gene expression counts of each sample were calculated by featureCounts (Liao *et al*., 2014). To get Nodal response genes, Nodal injected explants (different concentration 2 pg, 6 pg, 10 pg) were compared with WT explants at each timepoint, differential expressed genes were identified using DESeq2 package (Love *et al*., 2014). More than 4700 up regulated genes were identified (all up-regulated in Nodal injected explants) with pvalue <= 0.025 and log2FoldChange >= 0.75. To get Nodal immediate early genes, we first compared samples injected with 10 pg Nodal mRNA (treated with Cycloheximide or not) to WT (control) samples and got the differential expressed genes. We selected the genes with padj <= 0.1, pvalue <= 0.05, log2 fold change >= 1 (up-regulated) in each pair of differential expression analysis. Overlapped up-regulated genes were selected. 105 Nodal targets were identified in three timepoints (Supplementary Fig. S3N). DESeq2 was used to normalize the bulk RNA-seq data, and the normalized expression matrix of 105 Nodal targets in 36 bulk RNA-seq samples (except treated with Cycloheximide samples) was obtained. 105 Nodal targets were clustered into 4 groups using K-means algorithm. The heatmap was constructed by ComplexHeatmap (Gu *et al*., 2016). GO term enrichment analysis was performed by clusterProfiler (Yu *et al*., 2012) and top 8 GO terms (biological processes) were shown (Fig. 2C). cnetplot was performed by cnetplot function in clusterProfiler and top 10 GO terms were shown (Fig. 5E). Protein to protein interaction network (Supplementary Fig. S3O) was constructed by STRING (Szklarczyk *et al*., 2019) and visualized by Cytoscape (Shannon *et al*., 2003).

### Processing single-cell RNA-seq data

Illumina sequencing reads were aligned to the zebrafish mRNA reference genome (GRCz11 or GRCz10) using the 10x Genomics CellRanger pipeline (version 2.1.1) with default parameters. For the 4 hpf explants data, 4,303 cells were obtained; the median read depth per cell was 100,816 reads; the median genes per cell were 1,670 genes. For the 5 hpf explants data, 6,403 cells were obtained; the median read depth per cell was 68,301 reads; the median genes per cell were 1,321 genes. For the 6 hpf explants data, 8,364 cells were obtained; the median read depth per cell was 47,655 reads; the median genes per cell were 1,516 genes. For the 8 hpf explants data, 6,099 cells were obtained; the median read depth per cell was 78,963 reads; the median genes per cell were 2,167 genes. For the 10 hpf explants data, 10,760 cells were obtained; the median read depth per cell was 48,285 reads; the median genes per cell were 2,016 genes. For the 24 hpf explants data, 7,323 (9,399 GRCz10) cells were obtained; the median read depth per cell was 69,741 (54,337 GRCz10) reads; the median genes per cell were 2,077 (1,903 GRCz10) genes. The expressing matrix of each sample was obtained from CellRanger pipeline analysis and was used for next step analysis.

### Gene/cell filtering and cell types clustering

R package Seurat (version 2.3.4 or version 3.1.0) (Butler *et al*., 2018; Stuart *et al*., 2019) was used for cell clustering and data exploration. Initially, we created a Seurat object with following parameters: min.cells = 3, min.genes = 100. Next, cells were filtered using FilterCells function with appropriate parameters for each sample. NormalizeData function was used to normalize gene expressions of each cell and obtained the log-transformed expression counts. Highly variable genes were identified using FindVariableGenes function and top 2000 highest variable genes were selected for dimensionality reduction via principal component analysis (PCA). Appropriate PCs were used as input for clustering analysis (FindClusters function) after running JackStraw function and ScoreJackStraw function. We set different resolution parameters between 0.4 and 2 in FindClusters function, and certain cluster numbers were determined by distinguishing differential genes among clusters. FindAllMarkers function was used to find differentially expressed markers of each cluster. The clusters were visualized in two dimensions using *t*-SNE or UMAP. Finally, clusters were annotated according to ZFIN (Zebrafish Information Network) or other published studies (Farrell & Wang, 2018; Wagner *et al*., 2018).

### Dataset Integration

We integrated the Nodal injected explants datasets with zebrafish embryonic datasets at 24 hpf (Wagner *et al*., 2018) using a recently developed computational strategy from Seurat (Stuart *et al*., 2019). Firstly, individual datasets were preprocessed (log-normalization, data scaling for each gene, and detecting variable genes) as previous described with default parameters. Top 4000 genes were selected with highest dispersion, FindIntegrationAnchors function was used for canonical correlation analysis using 30 canonical correlation vectors. The anchors were used as input for IntegrateData function to integrate the two datasets together. Finally, integrated datasets were clustered into 33 groups using FindClusters function (resolution: 0.6) and FindConservedMarkers function was used to find conserved differentially expressed genes of each cluster.

### Single-cell trajectory analysis

Firstly, we constructed the Seurat object for the single cell data from embryos and Nodal injected explants. And then, we selected the overlapped genes (can be found in all single cell datasets, more than 17000 genes) for downstream analysis. We integrated the embryos data with Nodal injected explants data by MergeSeurat function from Seurat (version 2.3.4). The combined Seurat object was converted to h5ad format and used for scanpy (Wolf *et al*., 2018) analysis. Single cell trajectory was constructed following the steps suggested by scanpy, including total count normalization, log1p logarithmization and cell filtering. The batch effect correction was performed by this guidance (Luecken & Theis, 2019). Highly variable genes were identified by sc.pl.highly_variable_genes function. PCA was performed with n_comps = 50. A neighborhood graph among data points was computed and UMAP was used to generate a topologically faithful embedding with default parameters. Finally, PAGA (Wolf *et al*., 2019) was performed with default parameters and the layout ‘fa’ was used to construct the trajectory.

### Subclustering of *ndr2* positive cells

We focus on the cells that regulated by high Nodal signaling, *ndr2* positive cells were selected from the whole datasets (from 4 hpf to 10 hpf). Cell clustering and annotating were processed as previously described. The *ndr2* positive datasets were used for constructing URD developmental trajectories.

### Construction of a pseudospatial axis

UMAP can provide meaningful organization of cell clusters (Becht & McInnes, 2018). Firstly, we performed UMAP analysis for the whole cells or *ndr2* positive cells (4 hpf, 5 hpf and 6 hpf) of Nodal injected explants. The two dimensional embeddings from UMAP analysis used as input for Linear Discriminant Analysis (LDA) (https://CRAN.R-project.org/package=MASS). The clustering information was obtained from Seurat analysis as previous performed. The most significant dimensional embeddings after LDA dimension reduction (LD1) were used for the pseudospatial axis construction. The embeddings (LD1) were used for X-axis and random values (from −1 to 1) generated from runif function (R, https://www.r-project.org) were used for Y-axis. Cells were mapped to pseudospatial axis and labeled by different cell types. We explored the expressing cascade of Nodal targets along the pseudospatial axis using modified geneCascadeProcess from URD. Pseudospatial axis (X-axis) replaced pseudotime as the input for modified geneCascadeProcess function. We used 1000 genes random selected from all genes in Seurat object except Nodal targets and highly variable genes to calculate background expressing noise. At last, we used impulse model to fit genes’ expression in each segmented window.

### Construction of developmental trajectories

URD (Farrell & Wang, 2018) was used to reconstruct the developmental trajectories for *ndr2* positive cells (from 4 hpf to 10 hpf). The analysis procedure was performed as previous described (Farrell & Wang, 2018). We assigned early timepoint cells (4 hpf) as the root and the latest timepoint cells (10 hpf) as tips. Tip clusters were annotated by differential expressed markers in each cluster and EVL cells were removed before constructing the branching trajectory tree. The buildTree function from URD was used to construct the branching tree with following parameters: save.all.breakpoint.info = T, divergence.method = “preference”, cells.per.pseudotime.bin = 30, bins.per.pseudotime.window = 5, p.thresh = 0.001. cellsAlongLineage function was used to extract all cells along each lineage trajectory. geneCascadeProcess function was used to smooth genes’ expression using a moving window average (moving.window=5, cells.per.window=15) along each trajectory. The smoothed expression of each gene was fitted with an impulse model.

### Branchpoint Preference Plot of PP and endoderm

The branchpoint preference plot of PP and endoderm was constructed by URD with default parameters. Firstly, branchpointPreferenceLayout function was used to define the preference layout for the branchpoint. And then, stage, pseudotime information and gene expression were plotted on the branchpoint using plotBranchpoint function.

### Live imaging

Explant was placed in a glass bottom culture dish (Cellvis, D35-10-0-N) and was properly oriented in one drop of 0.1% agarose with the culture medium as previous described (Xu *et al*., 2014). And couple minutes later, the explant was covered with the culture medium. The explant live-imaged under Nikon CSU-W1 confocal microscope.

### Whole-mount in situ hybridization (WISH)

Whole-mount in situ hybridization was performed as described before (Thisse & Thisse, 2014). The constructs for *tbxta, sox32, sox17, chrd, noto pax2a, egr2b* and *gsc* were reported previously (Alexander & Stainier, 1999; Kikuchi *et al*., 2001; Krauss *et al*., 1991; Miller-Bertoglio *et al*., 1997; Oxtoby & Jowett, 1993; Schulte-Merker *et al*., 1994; Schulte-Merker *et al*., 1992; Talbot *et al*., 1995).To generate constructs of genes in Supplementary Table 1, a fragment of each cDNA was cloned into pEASY Blunt Zero Cloning vector(TransGen,CB501).The primers are listed in Supplementary Table 1. For single WISH, probes were labeled with digoxigenin-11-UTP (Roche Diagnostics), and substrate was NBT/BCIP. For double WISH, staining of embryos simultaneously hybridized with one probe labeled with digoxigenin-11-UTP and the other with fluorescein was performed using NBT/BCIP and fast red as substrates successively.

### Induction of secondary axis

Ventral overexpression of Nodal targets was done to induce secondary axis. For this, one random blastomere at the margin of the 16-32 cell stage dechorionated embryos was injected with approximately 4 pl mixture composed of specific mRNA and dye (rhodamine or fluorescein). Injected embryos that have fluorescence restricted to the ventral side were picked out at 6 hpf and incubated until 12 hpf, when embryos were fixed by 4% PFA for ensuing WISH analysis or imaged using a Fluorescent stereoscopic microscope (LEICA M205 FCA). 10 pg of *gsc* mRNA and 10 pg of *ndr2* mRNA were injected. 24 pg of ripply1 mRNA which is 7 times the molar amount of *gsc* was injected. All other prechordal plate specific Nodal targets were injected with the same molar amount as *ripply1* except *cxcl12a* (7 times the molar amount of *ripply1*).

## Data Access

The accession number for the RNA sequencing data reported in this paper is GEO: GSE165654. (https://www.ncbi.nlm.nih.gov/geo/query/acc.cgi?acc=GSE165654)

## Acknowledgments

We thank Dr. Bernard Thisse and Dr. Christine Thisse at University of Virginia, Dr. Jun Ma, Dr. Xiao-hang Yang, Dr. Feng He, Dr. Hong-qing Liang, Dr. Wan-zhong Ge and the members of Laboratory of Development and Organogenesis at Zhejiang University for helpful suggestions and discussions. This work was supported by grants from the Chinese National Key Research and Development Project (2018YFC1003203, 2019YFA0802402) and the National Scientific Foundation of China (32050109, 31970757, 31900576).

## Author Contributions

PFX and S.G.M designed research; YYX, TC, YFL, CL, YH and YJZ performed research; TC analyzed bulk RNA-seq and single cell RNA-seq data; TC, YYX, S.G.M and PFX wrote the manuscript; all authors reviewed and approved the manuscript.

